# Toward a genetic system in the marine cyanobacterium *Prochlorococcus*

**DOI:** 10.1101/820027

**Authors:** Raphaël Laurenceau, Christina Bliem, Marcia S. Osburne, Jamie W. Becker, Steven J. Biller, Andres Cubillos-Ruiz, Sallie W. Chisholm

## Abstract

As the smallest and most abundant primary producer in the oceans, the cyanobacterium *Prochlorococcus* is of interest to diverse branches of science. For the past 30 years, research on this minimal phototroph has led to a growing understanding of biological organization across multiple scales, from the genome to the global ocean ecosystem. Progress in understanding drivers of its diversity and ecology, as well as molecular mechanisms underpinning its streamlined simplicity, has been hampered by the inability to manipulate these cells genetically.

Multiple attempts have been made to develop an efficient genetic transformation method for *Prochlorococcus* over the years; all have been unsuccessful to date, despite some success with their close relative, *Synechococcus*. To avoid the pursuit of unproductive paths, we report here what has not worked in our hands, as well as our progress developing a method to screen the most efficient electroporation parameters for optimal DNA delivery into *Prochlorococcus* cells. We also report a novel protocol for obtaining axenic colonies and a new method for differentiating live and dead cells. The electroporation method can be used to optimize DNA delivery into any bacterium, making it a useful tool for advancing transformation systems in other genetically recalcitrant microorganisms.

## INTRODUCTION

While environmental microbiology has been revolutionized by the rapid pace of improved sequencing technologies [1], the number of genetically tractable model organisms has lagged behind [2]. The dearth of such organisms has limited our progress since most ‘omics’ analyses rely on comparisons with model organisms for their interpretation [1], [3]–[5]. Isolation of pure (axenic) cultures from the wild has proved to be a significant challenge and developing genetic tools for these isolates has been even more difficult.

Countless attempts have demonstrated that the creation of successful genetic transformation protocols requires tedious trial and error, and methods developed for one strain are often unsuccessful in close relatives (for a few examples see: [6]–[13]). There is no single solution for all bacteria, and no way to predict which techniques will succeed [14]. Thus, there is a pressing need for broad-spectrum, automatized, and standardized genetic transformation procedures [15] that could enable downstream genetic screening methods such as transposon insertion sequencing [5], [16]–[18]. Such projects are not suitable for graduate students or post-docs because of the high risk involved, nor are they easily funded.

*Prochlorococcus* is the most abundant cyanobacterium worldwide, dominating vast regions of the global oceans [19]. It may account for up to 50% of the chlorophyll in oligotrophic ocean regions [20], [21] and is responsible for around 8.5% of global ocean primary productivity [19]. Our understanding of *Prochlorococcus* biology and ecology has been greatly facilitated by the rise of metagenomic, metatranscriptomic, and metaproteomic studies of ocean samples over the past decade [22]–[26]. Because of its high relative abundance, it often dominates databases derived from surface microbial communities in the oceans [27], and has emerged as one of the best-described marine microorganisms with more than one thousand complete or nearly complete genomes available [28]–[30]. *Prochlorococcus* is a perfect example of the imbalance between the availability of genomic data and the dearth of genetic tools.

*Prochlorococcus’* numerical dominance in oligotrophic oceans is attributed to its small size (hence high surface/volume ratio), which enhances its ability to compete for limiting nutrients [31], and its vast genomic diversity [32]–[37], which expands the niche dimensions of the *Prochlorococcus* meta-population [28], [38], [39]. Ecologically meaningful units within the meta-population can be found at all levels of phylogeny, both deeply rooted – such as their adaptation to different light levels – and in the “leaves of the tree” where differences in the presence/absence of specific nutrient acquisition genes can be found [28], [37], [38], [40]. Although we have been able to unravel the selective pressures that shape the distributions of genes of obvious ecological relevance, many intriguing stories are undoubtedly obscured by our inability to assign functions to the numerous unannotated genes in these cells; for each new *Prochlorococcus* strain isolated, or wild single-cell sequenced, roughly 100 new genes are added to the *Prochlorococcus* pangenome – the vast majority of which are of unknown function [28].

In addition to its ecological and biogeochemical relevance, the biology of *Prochlorococcus* is of interest because of its potential as a chassis for synthetic biology applications [41]. This cell is the simplest photosynthetic “machine” designed by nature – an attractive foundation for manipulation [41]–[44]. It has the smallest genome of any oxygenic phototroph as well as an efficient carbon concentrating mechanism [45]. Indeed, a recent study screening the metabolic potential of cyanobacteria for biofuel production ranked various *Prochlorococcus* strains as the top candidates [46]. Furthermore, their diversity makes them ideal subjects for studying the properties of small-genome organism consortia for bioproduction [43].

Progress in studying *Prochlorococcus* from these different perspectives has been hampered by our inability to develop a robust genetic system. Although we and others have worked on this over many years, very limited progress has been made. Possible reasons behind these failures include the cell’s slow growth rate (roughly one doubling per day under optimal laboratory conditions), their reluctance to grow axenically on solid media, specific media requirements [47], [48], sensitivity to trace metal contamination [49], [50] and reactive oxygen species [51], unusual membrane composition [52], and their apparent reduced homologous recombination potential [53].

From our perspective, a significant impediment to rapid progress is that investigators generally do not publish negative results or partial progress; as a result, considerable time is wasted by others in rediscovering what does not work. Thus, before describing our progress, we first report experiments that failed. We initially tried to build upon the only reported successful (albeit inefficient) *Prochlorococcus* genetic transformation method, based on conjugation from an *E. coli* donor strain [54]. This method proved unsuccessful in our hands, probably because the 6-month multi-step procedure is vulnerable to user variability. From there we moved on to explore the use of electroporation and were able to find appropriate conditions for delivering DNA into living *Prochlorococcus* cells – developing a method that should be applicable to a wide range of microorganisms. The next step will be to develop genetic tools compatible with the host that will also bypass the cell’s defences against exogenous DNA, which are abundant in cyanobacteria [55]. We report the results of some initial attempts, in which transposomes were delivered into *Prochlorococcus* cells.

## RESULTS AND DISCUSSION

### Selecting an antibiotic for *Prochlorococcus*

Finding a suitable selectable marker is a prerequisite for the selection of transformants. We subjected different *Prochlorococcus* strains to various common antibiotics in order to find their minimum inhibitory concentration (MIC), revealing that some antibiotics were more effective than others (Table 1). While *Prochlorococcus* strain MED4 was resistant to nalidixic acid, it was highly sensitive to ciprofloxacin and chloramphenicol (Table 1). Kanamycin, which has been used as a selective marker for various marine *Synechococcus* strains [6] and a LLIV clade *Prochlorococcus* strain [54] had a MIC of 50 μg ml^−1^, and failed to suppress the emergence of spontaneous kanamycin-resistant colonies at a high frequency in our hands, regardless of the *Prochlorococcus* strain. We ultimately focused on streptomycin, which can be used in combination with spectinomycin to reduce the appearance of spontaneous resistance mutants [56], [57].

**Table 1.**
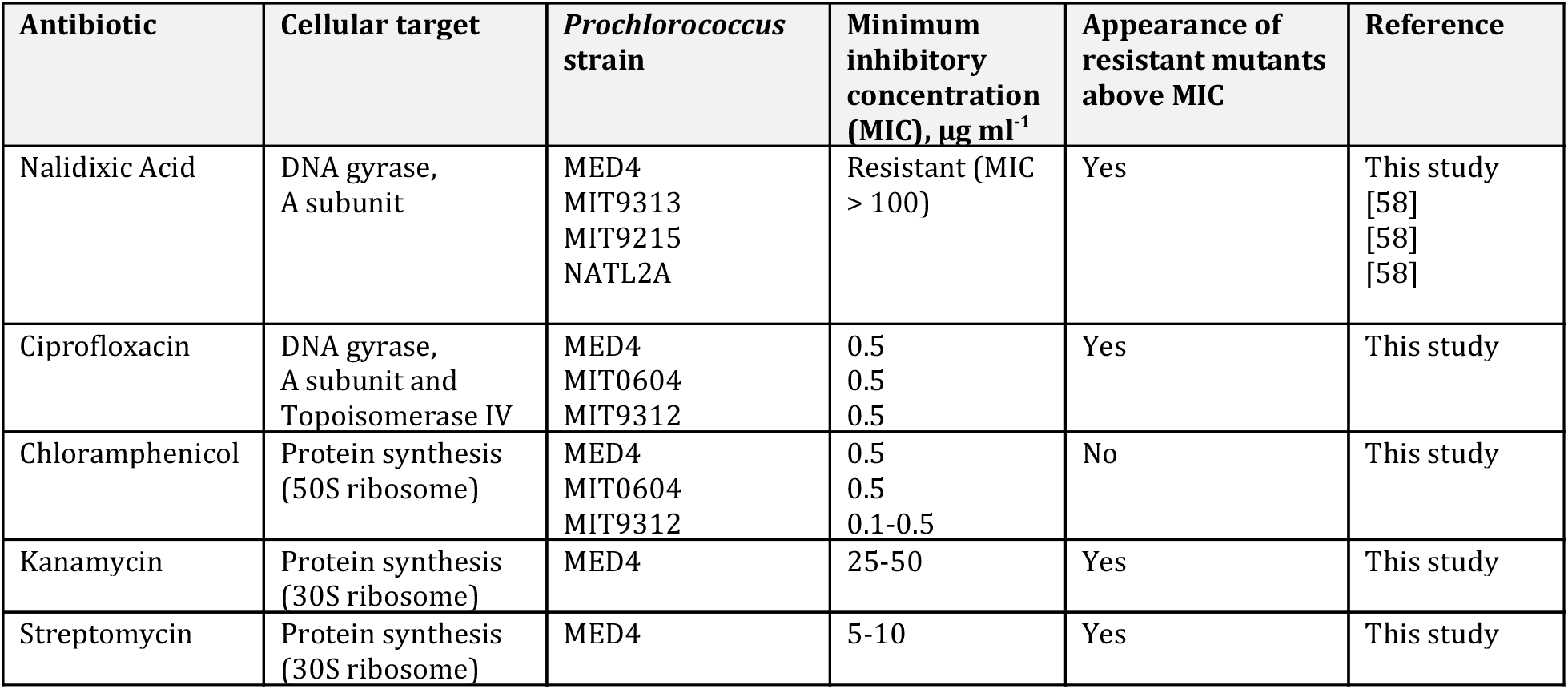
Minimum inhibitory concentration (MIC) of various antibiotics on different *Prochlorococcus* strains.

| Antibiotic | Cellular target | *Prochlorococcus* strain | Minimum inhibitory concentration (MIC), $\mu\text{g ml}^{-1}$ | Appearance of resistant mutants above MIC | Reference |
| --- | --- | --- | --- | --- | --- |
| Nalidixic Acid | DNA gyrase, A subunit | MED4<br>MIT9313<br>MIT9215<br>NATL2A | Resistant (MIC > 100) | Yes | This study<br>[58]<br>[58]<br>[58] |
| Ciprofloxacin | DNA gyrase, A subunit and Topoisomerase IV | MED4<br>MIT0604<br>MIT9312 | 0.5<br>0.5<br>0.5 | Yes | This study |
| Chloramphenicol | Protein synthesis (50S ribosome) | MED4<br>MIT0604<br>MIT9312 | 0.5<br>0.5<br>0.1-0.5 | No | This study |
| Kanamycin | Protein synthesis (30S ribosome) | MED4 | 25-50 | Yes | This study |
| Streptomycin | Protein synthesis (30S ribosome) | MED4 | 5-10 | Yes | This study |

### Attempts at *E. coli* mediated conjugation

*E. coli* mediated conjugation has been successfully used to transform several marine *Synechococcus* strains (which are closely related to *Prochlorococcus*) [6], [59] and has been reported to work for one LLIV clade *Prochlorococcus* strain [54]. We first attempted *E. coli* conjugation with *Prochlorococcus* via a filter mating procedure which did not yield any conjugants in our hands, even after incubating for 2 months. We next used a liquid mating procedure, testing different *E. coli* donor strains carrying plasmids with distinct antibiotic resistance genes on various *Prochlorococcus* and *Synechococcus* recipient strains (see methods for details, and strains and plasmids in Table 2), but again obtained no conjugants.

**Table 2.**
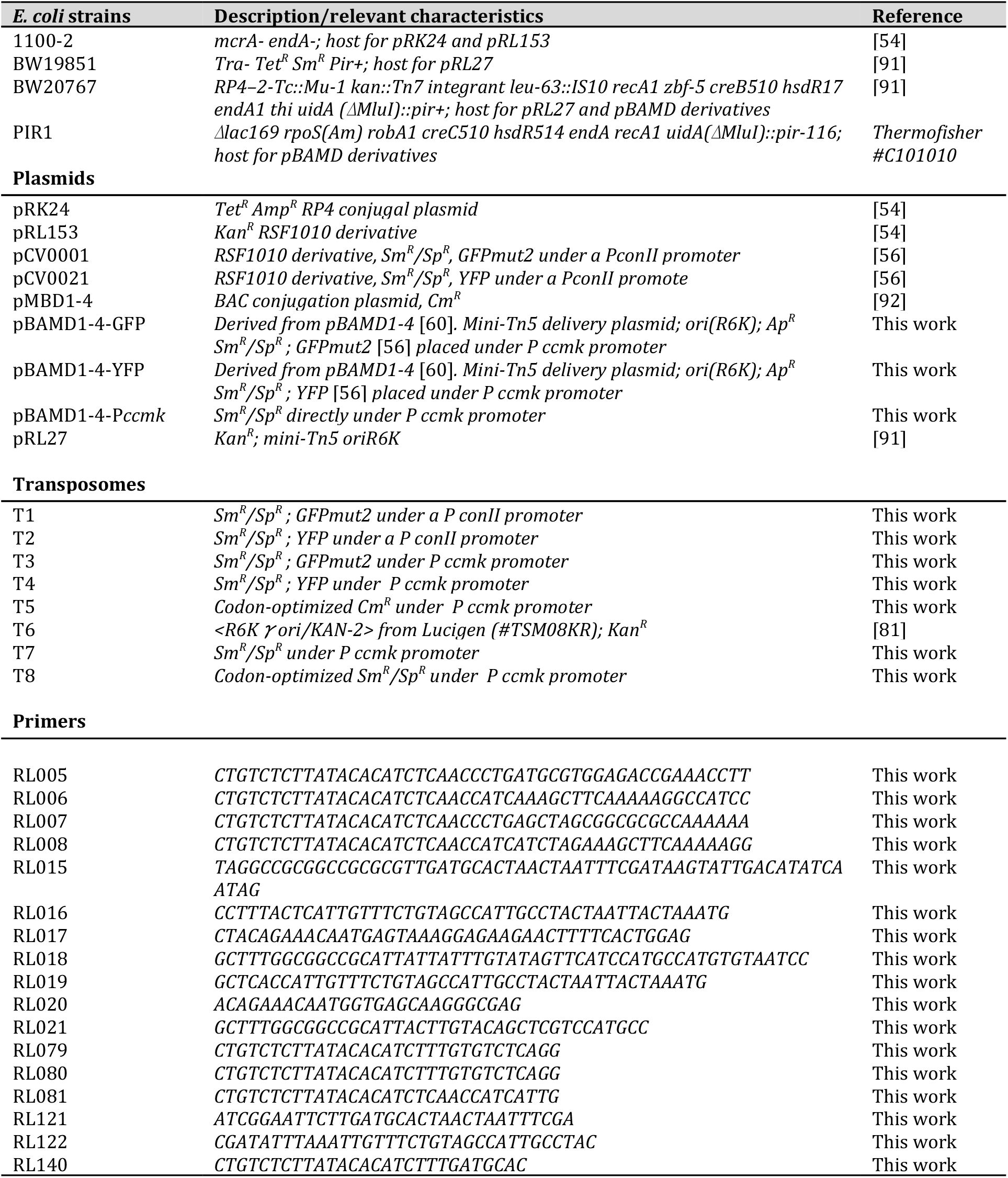
Strains, plasmids and primers. Antibiotic markers: Ap^R^, ampicillin; Cm^R^, chloramphenicol; Gm^R^, gentamicin; Km^R^, kanamycin; Sm^R^, streptomycin; Sp^R^, spectinomycin; and Tet^R^, tetracycline.

One of the limitations of our mating experiments was the low survival frequency of *Prochlorococcus* during filter mating. Indeed, while a *Synechococcus* strain could readily recover after the mating procedure, *Prochlorococcus* strains were often lost at this step. The switch to a liquid mating procedure (see methods) improved survivability but still did not yield transconjugants. In an effort to find an alternative route to conjugation with *Prochlorococcus*, we developed a mixed approach where mating was performed inside an agar stab (see methods for details). Briefly, concentrated cultures of the donor and receiver strains were mixed and injected together inside 1 mL of polymerized agar - the ‘agar stab’ (see supplementary Fig. 1). We reasoned that this agar stab environment would provide a solid medium favoring conjugation while mitigating the drawback of desiccation on top of a filter. After 24h incubation, the cells were removed from the agar with a pipet tip, resuspended in liquid medium, and plated on antibiotic selection plates.

We attempted this procedure with conjugative plasmid - pBAMD1-4 [60] - containing a resistance gene to spectinomycin/streptomycin. Plasmid pBAMD1-4 is a suicide vector containing the mini-Tn5 transposition system. This transposition system, encoded on vector pRL27, was previously used successfully in *Synechococcus* strain WH8102 [61] and by Tolonen et al. in *Prochlorococcus* strain MIT9313 [54]. Though this protocol could not be reproduced in our hands, it shows that Tn5 transposition can function in *Prochlorococcus*. In addition, we cloned the *Prochlorococcus* promoter for Rubisco protein, P *ccmk*, in front of a fluorescent reporter *gfp* or *yfp* gene to facilitate the characterization of conjugants (see methods for details). Using this ‘agar stab’ procedure, a *Synechococcus* strain and two *Prochlorococcus* strains survived and were able to grow on plates in the absence of antibiotics. After 30 days, we obtained transconjugant colonies of *Synechococcus* WH8102, but did not obtain transconjugants for either *Prochlorococcus* strain (SB and NATL2A). We thus abandoned conjugation as a feasible mechanism of genetic transfer and moved on to electroporation. This required the development of a method to assess DNA delivery into live cells.

### Development of a dead cell stain for *Prochlorococcus*

In the absence of an established method to transform *Prochlorococcus*, we decided to focus on optimizing the initial step required for any genetic system: DNA delivery. A screening procedure for DNA entry must be able to differentiate between live and dead cells since the delivery of DNA inside dead cells would yield false positives. We needed a dead cell stain that emited in the blue/violet wavelength range (between 400-500 nm wavelength) so that it could be differentiated from chlorophyll autofluorescence (650-700 nm), and fluorescein-labelled oligonucleotide fluorescence (495-555 nm). While Sytox green (ThermoFisher Scientific) has been used successfully with *Prochlorococcus* [51], we found that under certain conditions, such as cells recovering from stress events, it does not effectively distinguish live from dead cells [62]. We next tried Sytox blue (ThermoFisher Scientific), which emits around 480 nm, but the signal was below the detection limit of the Guava easyCyte HT flow cytometer. We then turned to an amine reactive stain – the Live/Dead™ fixable Violet stain (ThermoFisher Scientific), which was developed primarily for eukaryotic cells and emits fluorescence at 452 nm. This stain reacts with amine residues from proteins; it labels the exposed amine groups at the surface of both live and dead cells, but only dead cells that allow the dye to penetrate their membrane reveal labelling in the cytoplasm, greatly increasing the overall signal. This stain identified dead *Prochlorococcus* cells in a robust and highly reproducible way (Fig. 1), representing a new addition to the *Prochlorococcus* tool kit.

**Fig. 1.**
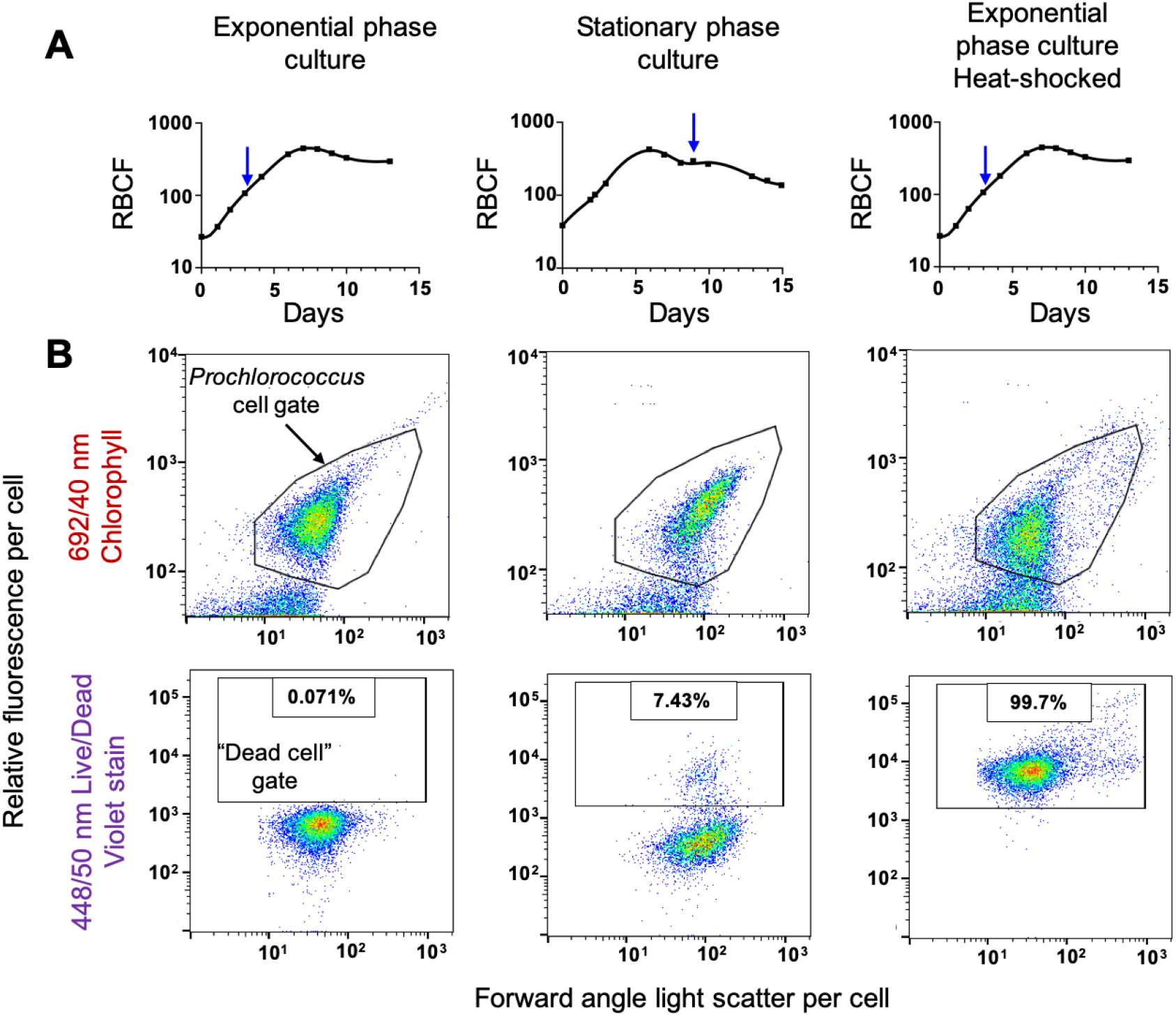
Efficacy of Live/Dead™ stain in differentiating live and dead *Prochlorococcus* cells. **A)** Growth of the culture as measured by bulk relative red fluorescence as a function of time. Blue arrows indicate when samples were taken for treatment and flow cytometric analysis. The third growth curve indicates when, in the growth curve, a sub-sample was taken for the heat-shock measurement. RBCF: Relative Bulk Chlorophyll Fluorescence. **B)** The gates in the upper flow cytometry panels delineate the *Prochlorococcus* cell population, while the rectangular gates in the lower plots indicate the increase in the number of dead cells as the culture progresses from exponential to stationary phase culture, and also after heat shock (bottom right). Percentages of dead cells measured in each population are indicated.

### Development of an efficient electroporation procedure

While electroporation is one of the most powerful techniques for delivering DNA to the inside of living cells [14], [63], and protocols have been developed for marine bacteria [64]–[67], it is a harsh treatment, typically resulting in significant cell loss [68]. Further, the required removal of salts results in osmotic stress - particularly challenging for a notoriously temperamental marine bacterium like *Prochlorococcus* [47]. This stress must be mitigated using an osmoprotectant, and we explored the efficacy of different osmoprotectants as we pursued this approach.

#### Selecting an osmoprotectant

We tested the ability of *Prochlorococcus* to survive exposure to most commonly used electroporation osmoprotectants (glycerol, sorbitol, and PEG8000, see methods for details). Cells washed in glycerol or PEG8000 did not survive any better than cells washed with MilliQ water, while sorbitol allowed cells to recover as fast as the seawater media control (Fig. 2). Of note, the initial cell density had a major impact on recovery: a starting density of 3.3 × 10^7^ cells μl^−1^ yielded only 0-5% recovery, whereas one of 3.7 × 10^8^ cells μl^−1^ yielded 53 ± 4% recovery, possibly due to more efficient pelleting or mitigation of oxidative stress at higher densities. Washes with sorbitol at the higher densities, however, yielded sufficient cell numbers for electroporation.

**Fig. 2.**
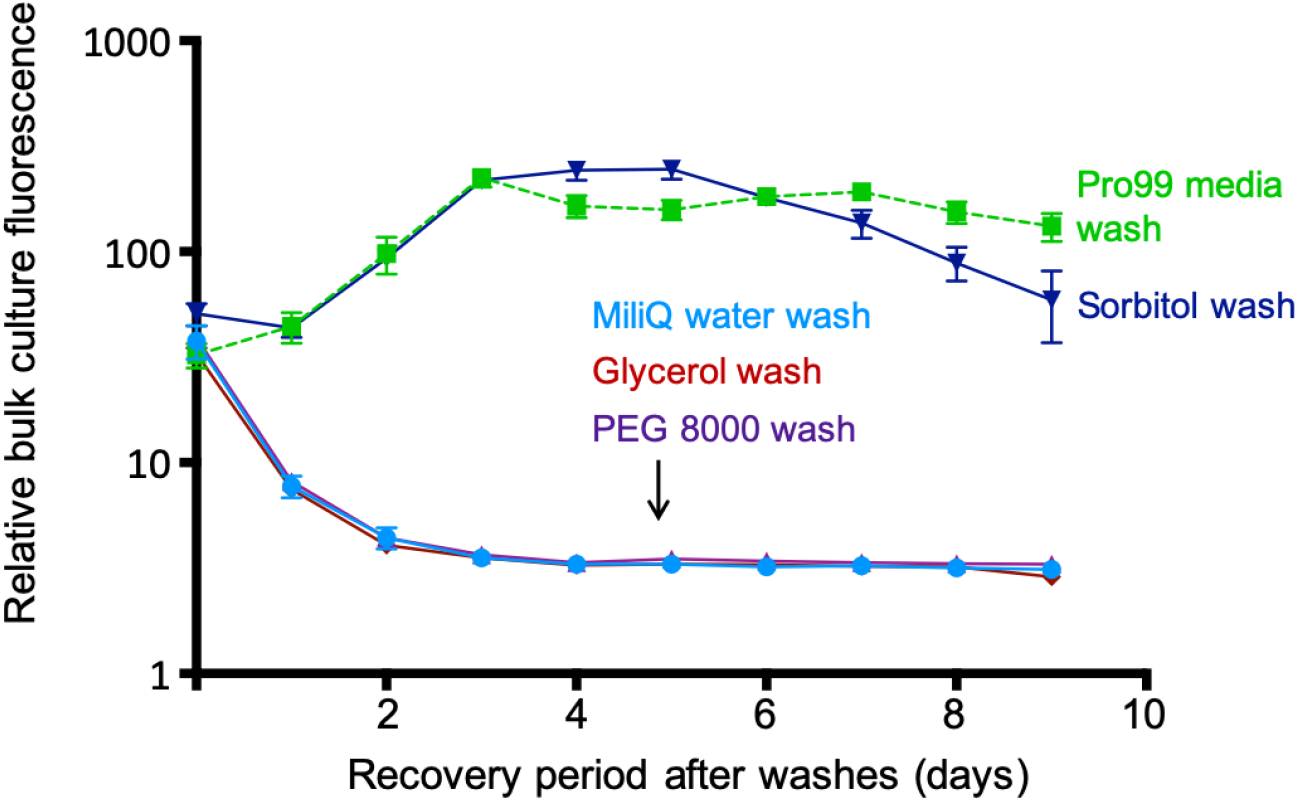
Effects of different osmoprotectants on the survival of *Prochlorococcus*. *Prochlorococcus* strain MED4 cells were washed three times in glycerol 10% (v/v), sorbitol 18.2 % (w/v) (corresponds to 1 M), and PEG 8000 5% (w/v), placed back in their growth medium (Pro99), and culture growth was monitored using bulk fluorescence. Washes were also performed without any protectant in MilliQ water as a negative control, and in the growth medium Pro99 as a positive control. Error bars show the standard deviation of triplicate samples.

#### Electroporation optimization

Using sorbitol as the osmoprotectant buffer, we next set up a method to screen for the most efficient electroporation conditions. Briefly, exponentially growing cultures were harvested and suspended in sorbitol. Fluorescein-labelled DNA was added to the washed cells, which were electroporated at variable electric fields and time constants while keeping the capacitance and resistance fixed. Cells were then transferred to fresh media for recovery and were used for dead cell staining and flow cytometry analysis (see Fig. 3A and material and methods section for details). We initially attempted to deliver a fluorescein-labelled plasmid in these optimization experiments; however, *Prochlorococcus* exhibits strong autofluorescence in the 650-700 nm range due to chlorophyll [69], [70], but also a lower background autofluorescence in the green range used to detect fluorescein. This residual autofluorescence was sufficient to mask the weak signal obtained from fluorescein-labelled plasmids and prevented their reproducible detection. We thus used fluorescein-labelled oligonucleotides as our probe for DNA delivery (see methods for detail). Oligonucleotides can be delivered in much higher amounts than plasmids, allowing us to increase the signal to noise ratio significantly. Although oligonucleotide delivery cannot predict plasmid delivery quantitatively - especially considering that plasmid size is inversely correlated with delivery efficiency [71] - their identical molecular composition allows them to be a convenient proxy to assess the best conditions for delivery of DNA molecules by electroporation.

**Fig. 3.**
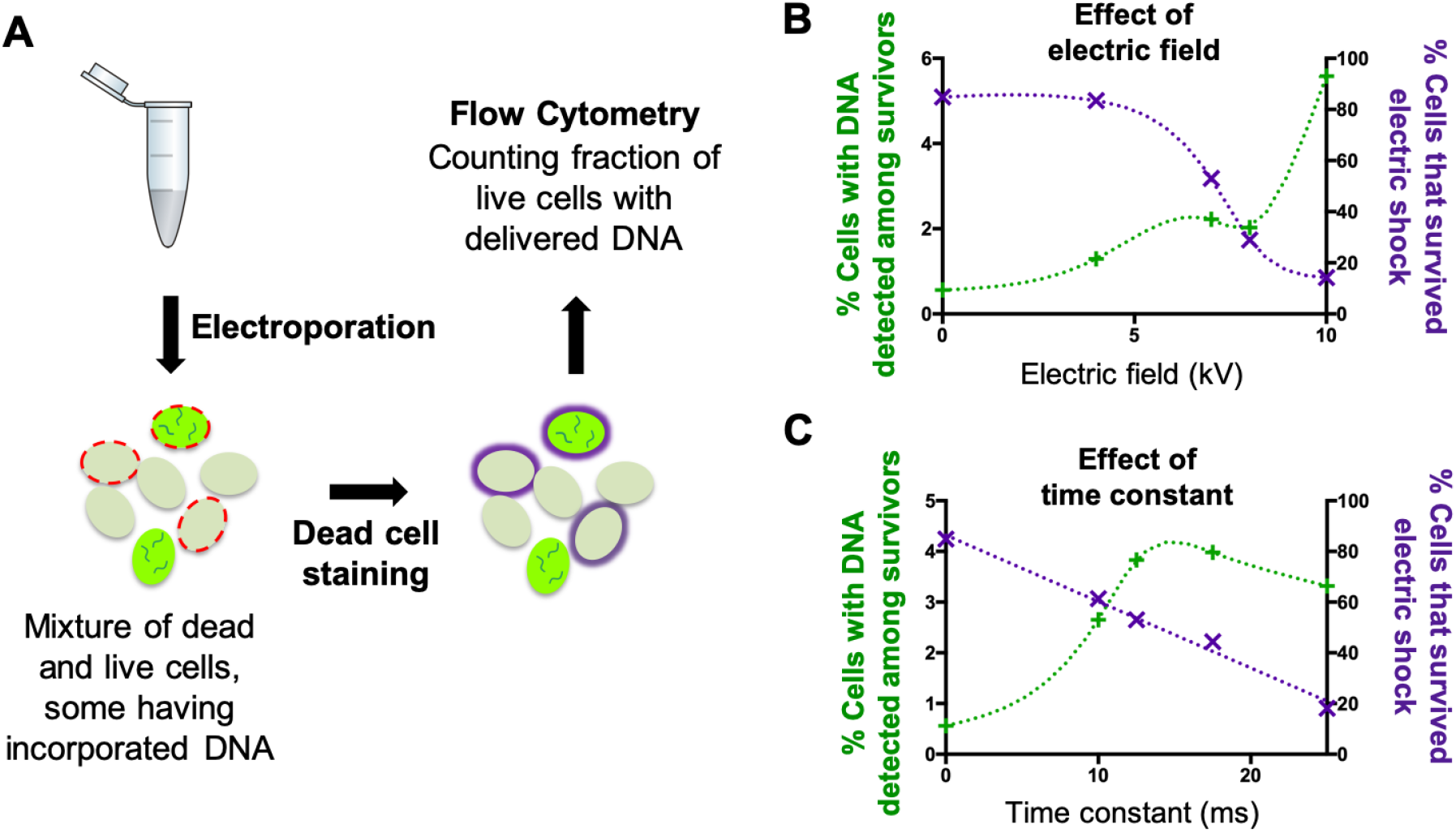
Screening workflow for examining the efficiency of DNA delivery into cells via electroporation. **A)** DNA delivery screening workflow. Light green oval shapes represent *Prochlorococcus* cells; oval shapes filled with bright green and short wavy lines correspond to cells that have incorporated the fluorescently labelled oligonucleotides; red dotted lines represent non-viable cells compromised by the electroporation treatment; solid purple line corresponds to non-viable cells stained with the dead-cell stain. **B)** Percentage of live cells (assessed by the Live/Dead violet stain) with detectable levels of fluorescein-labelled oligonucleotides as well as the % of cells surviving the electric shock, as a function of electric field intensity. **C)** Same as B but varying the time constant of the electroporation shock.

We next tested a wide range of electroporation conditions. Using a fixed time constant of 12.5 ms, we found electric fields of 7 kV/cm to be the most efficient condition for balancing the delivery of DNA inside cells while minimizing cell death (Fig. 3B). We then varied the time constant, and found a peak delivery rate at about 15 ms; cell mortality appeared to increase linearly with the time constant (Fig. 3C). This broad exploration of electric fields and time constants allowed us to find the most promising conditions for DNA delivery using a minimum number of samples. Considering the two-week time frame needed to grow enough cell material, and the limited number of samples that can be tested in parallel, this approach proved particularly advantageous.

We next focused on further optimizing electroporation conditions over the narrow range between 7-8 kV/ cm (Fig. 4). Increasing the time constant from 12.5 ms to 25 ms at 7 kV/cm marginally increased the delivery rate, but decreased cell survival more than two-fold. On the other hand, using 8 kV/cm with a shorter time constant of 5 ms increased the number of survivors but reduced the DNA delivery rate to that of the non-electroporated control (Fig. 4A). Thus, taking into account both the DNA delivery efficiency and cell survival, we determined that 7 kV/ cm; 12.5 ms was the most efficient condition for electroporation (Fig. 4B). That such small variations in either the electric field or the time constant had dramatic effects was unexpected, and unusual for optimization of electroporation conditions [65], [72]–[74].

**Fig. 4.**
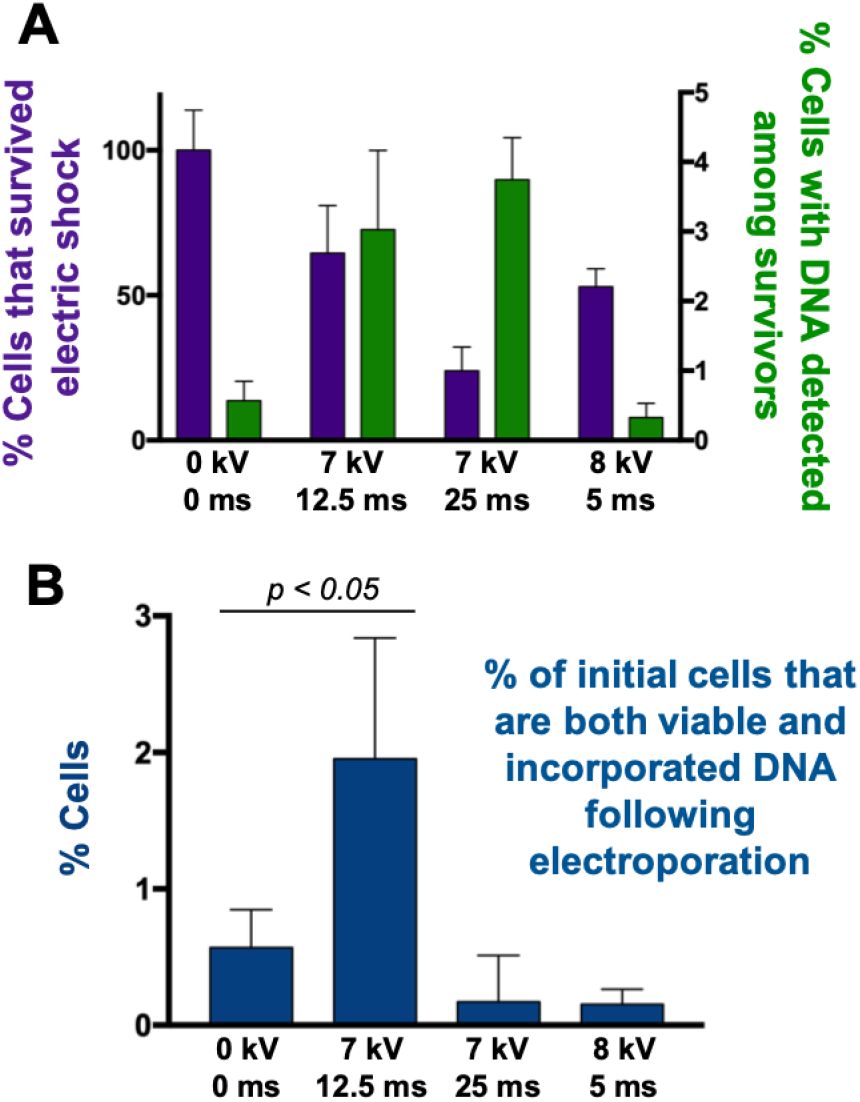
Fine scale optimization of electroporation conditions. **A)** Cell survival following electric pulse (assessed by dead-cell stain, left axis), and percentage of survivors with detectable levels of fluorescein-labelled oligonucleotides (right axis). **B)** Efficiency of DNA delivery into cells for these electroporation conditions. T-test p value is indicated for the best electroporation conditions compared to the control. Note that a single electroporation reaction is performed on approximately ~10^9^ cells; thus, 2% represents a large number of potential transformants.

### Development of a pour-plating procedure for axenic *Prochlorococcus* cultures

Assuming that mutants will one day be generated, we will need a procedure for selecting them on solid media. To date, the only way to grow *Prochlorococcus* on solid media has been to use a ‘helper strain’ of heterotrophic bacteria, typically *Alteromonas* [48], [51], [75]. *Prochlorococcus* does not encode catalase, thus the ‘helper’ strain provides this function – removing reactive oxygen species from the media, reducing oxidative stress, and thereby facilitating growth [76]. Since this approach then requires re-purification of axenic *Prochlorococcus* after obtaining a mutant, we sought to develop a plating protocol that would eliminate the helper cells and the need for this additional purification step. We approached this in two ways. First, because autoclaving agar produces reactive oxygen species [77], we sterilized the low melting point agar solution in ultrapure water using three rounds of boiling in a microwave before adding it to Pro99 medium resulting in a 0.3% agar solution. Second, we included pyruvate in the medium, which serves to quench reactive oxygen species and is known to enhance the survival of liquid axenic *Prochlorococcus* cultures at low cell densities [78], [79]. With these modifications, we were able to obtain pour plates containing axenic *Prochlorococcus* colonies (see Methods for details). The colonies typically took 2-3 months to become visible to the naked eye (Supplementary Fig. 2), and could then be picked and transferred into small (~5 mL) volumes of liquid Pro99 medium; they grew as expected and were verified as axenic. This combination of microwave sterilization of the agar and addition of pyruvate to the plating medium allowed us to reproducibly obtain axenic colonies for all the *Prochlorococcus* strains tested so far (MED4, MIT9313, MIT0604, MIT1314, MIT9312, and SB).

### Transformation attempts by transposome electroporation

The vast majority of genetic systems developed for bacteria rely on the use of replicative plasmids. Replicative plasmids offer a versatile platform to introduce and express exogenous genes in the target bacterial strain, allowing the development of further technologies for gene knockouts, such as Lambda Red recombineering or CRISPR-Cas9 [14]. However, in order to replicate, plasmids require an origin of replication compatible with the host. A common strategy for engineering a replicative plasmid is to use an origin of replication sequence derived from a wild plasmid found in the target bacterial host or to use a ‘broad host range’ origin of replication derived from conjugative plasmids [80]. As of yet, no wild plasmids have been found associated with *Prochlorococcus* cells, which are under intense selective pressure for genome minimization [28], [31]. Further, broad host range plasmids have yielded mediocre maintenance in closely related marine *Synechococcus* strains [56] and have failed to replicate at all in *Prochlorococcus* [54].

An alternative strategy is to use integrative plasmids to serve as delivery vectors, relying on the target cell’s homologous recombination machinery to recombine the exogenous DNA into the host genome [14]. This process has been successful in generating gene knockouts in marine *Synechococcus* strains [6] but required a lengthy procedure for *Prochlorococcus* [54], which was not reproducible in our hands (see section above on conjugation attempts). We thus opted for a strategy that relies minimally on the host’s genetic abilities - the use of transposase enzymes to integrate foreign DNA into the target host chromosome. The transposase gene and transposon DNA can either be introduced on a plasmid vector so that transposase expression and activity occurs within the target cells (for example, plasmid pBAMD14 [60] used in conjugation attempts), or the transposase/transposon protein-DNA complex (the transposome) can be prepared in vitro and delivered inside the host by electroporation [81].

We used the EZ-Tn5 transposition system (Lucigen), which has worked successfully for several challenging bacteria [82]–[85]. Transposon DNA inserts were designed using genetic segments optimized for cyanobacteria [56], the mini-Tn5-vector pBAMD [60], and sequences designed specifically for *Prochlorococcus*, such as the strong promoter P *ccmk* [86] or the codon-optimized *aadA* and *cat* resistance genes (see methods for details). Each of these constructs was tested in two *Prochlorococcus* strains (MED4 and MIT9313), as well as in several other strains selected for a lower likelihood of degrading incoming DNA by means of a restriction-modification system (MIT1314, MIT0604, MIT9215). However, no transformants were obtained in any of these attempts, despite growth of electroporated strains on the no-antiobiotic on control plates.

### Looking toward the future

While suitable DNA delivery conditions is a significant step forward, we have not yet achieved a successful genetic transformation of *Prochlorococcus*. Among the remaining hurdles are overcoming host defences against foreign genetic material such as restriction-modification systems, and optimizing expression of the antibiotic selection marker and the stable integration of foreign DNA inside the chromosome. We are confident, however, that effective solutions can be found to overcome these hurdles. Solutions might come from new transposome systems [87], which could reveal more efficient than the Tn5 transposition; from the delivery of Cpf1-RNA complexes inside cells [88], [89]; from using more systematic approaches to evade restriction-modification defences [15]; through the discovery of yet unknown defence systems [90] that are hampering transformation; or finally by harnessing endogenous mobile genetic elements, which have been tailored by evolution to work efficiently in these minimal cells. The latter is the direction we are currently exploring.

## MATERIALS AND METHODS

### Culture conditions

Axenic *Prochlorococcus* MED4 cells were grown under constant light flux (30–40 μmol photons m^−2^ s^−1^) at 24°C in natural seawater-based Pro99 medium containing 0.2-μm-filtered Sargasso Sea water, amended with Pro99 nutrients (N, P, and trace metals) prepared as previously described [47]. Growth was monitored using bulk culture fluorescence measured with a 10AU fluorometer (Turner Designs).

### Testing different osmoprotectants

For each osmoprotectant tested, two mL of triplicate exponentially growing MED4 culture were harvested by centrifugation at 13,000 g for 10 min. Under sterile conditions, pellets were resuspended by gently pipetting up and down in a volume of 100 μL of either solutions10% (v/v) glycerol in water; 18.2% (w/v) sorbitol in water (1M); 5% (w/v) PEG8000 in water; Pro99 medium (positive control); or MiliQ water (negative control). The washing was repeated once before cells were centrifuged again, resuspended into 1 mL of Pro99 medium, and inoculated in 5 mL of Pro99. The tubes were placed back in the incubator, and recovery was monitored by measuring bulk culture fluorescence. Of note, all osmoprotectant solutions were filter sterilized through 0.2 μm supor membrane filters. Cultures that recovered were checked for the presence of heterotrophic contamination using purity broths, as previously described [47] to ensure that recovery was not facilitated by a contaminating heterotrophic partner [75].

### Testing different antibiotics

MED4 cells were grown in Pro99 medium with a Sargasso seawater base, at a light level of 25 μmol photons m^−2^ s^−1^. Growth was monitored by measuring bulk chlorophyll fluorescence and was compared to a no-drug control culture.

### Strains, plasmids, and transposomes

### *E. coli* mediated conjugation

For conjugation via filter mating, we followed the procedure described by Tolonen et al [54]. Conjugation was performed using *E. coli* 1100-2 carrying plasmid pRK2 (as the conjugation vector) and pRL153 (kanamycin-resistant), as well as *E. coli* BW19851 carrying BAC conjugation plasmid pMBD14 [92] (chloramphenicol-resistant) - on receiver strain *Synechococcus* 8102, and *Prochlorococcus* strains MIT9313 and MED4. As described above, in our hands, the method did not yield any conjugants after incubation of exconjugants for 2 months. For conjugation via liquid mating, we used the same donor and receiver strains, and the following procedure: 1 mL of log-phase donor *E. coli* was added to 25 mL of log-phase *Synechococcus* or *Prochlorococcus* receiver strain and the co-culture was left to grow overnight. Antibiotic was then directly added to the co-culture at the same concentration used on solid media (kanamycin 50 μg ml^−1^, chloramphenicol 10 μg ml^−1^). The addition of nalidixic acid at 50 μg ml^−1^ was also attempted to remove the *E. coli* donor strain. Cultures were then grown for 45 – 60 days but did not yield conjugants.

For conjugation via “agar stab mating”, we used *E. coli* donor strain BW20767 carrying plasmids pBAMD1-4-GFP or pBAMD1-4-YFP for mating with *Synechococcus* WH8102 and *Prochlorococcus* strains NATL2A and SB. 25 mL of log-phase receiver strain was pelleted by centrifugation at 5000 g for 15 min at 20°C and resuspended in 200 μL Pro99 medium. Log phase *E. coli* donor strain was mixed at a 1:1 cell ratio with the receiver strain, and 20 μL of the mixture was injected inside a 1 mL volume of polymerized Pro99 with 1% agar - the ‘agar stab’ (see supplementary Fig. 1). After 24 h incubation, cells were sucked out of the agar using a pipet tip, resuspended in liquid Pro99 medium, and plated for selection with 5 μg ml^−1^ of spectinomycin and streptomycin (combined 1:1; plating procedure below).

### Electroporation procedure

Exponentially growing cultures were harvested by centrifugation at 7,000 g for 15 min at 20°C. 50 mL culture pellets were washed 3 times in 5 mL (1 M sorbitol solution) by centrifugation at 5000 g for 8 min at 20°C. The final resuspension volume was calculated so that cells were concentrated 1000 times compared to the initial culture volume. 500 ng of 90 bp fluorescein-labelled oligonucleotides (sequence: cctcataacaagcagcgctcatagtattaggaatatcgtgaaattcaagatctaagaatatttttttatttaaatttttcaaaattttta) were added to 50 μL of the washed cells and electroporated in 2 mm gap cuvettes at variable electric fields and time constants while keeping the capacitance and resistance fixed at 25 μF and 200 ohms. One mL of fresh Pro99 medium at room temperature was immediately added to the cuvette, and cells were transferred to a culture tube containing 4 mL Pro99 supplemented with 5 mM glucose and 5 mM pyruvate for recovery. Samples were then directly stained with Live/Dead violet and analyzed by flow cytometry. We initially tried to deliver a fluorescently labelled pUC19 plasmid (using the fluorescein LabelIT Tracker kit, MirusBio), but the signal to noise ratio was not sufficient to detect the plasmid.

For transposome delivery, 2 μL of EZ-Tn5 (Lucigen) *in vitro*-assembled transposomes were added to the 50 μL cell suspension in sorbitol before electroporation. Addition of DOTAP liposomal transfection reagent (Millipore Sigma) and Type One Inhibitor (Lucigen), following the manufacturer’s instructions, were also attempted, as they were shown to favour successful transposome transformation in the cyanobacterium *Arthrospira platensis C1* [93], [94]. After transposome delivery, cells were left to recover for 24 h before plating.

### Live/Dead violet Staining protocol

For each sample labelled, 100 μL of culture were transferred to an Eppendorf tube containing 900 μL of filtered Pro99 medium. 1 μL of reconstituted LIVE/DEAD™ Fixable Violet Dead Cell Stain (Thermo Fisher Scientific) was added to the 1 mL cell suspension and incubated for 30 min at room temperature in the dark. Cells were pelleted by centrifugation at 10,000 g for 5 min at 20°C and resuspended in 0.5 mL of fresh Pro99. Samples were then serially diluted and immediately run on the flow cytometer. A positive control (untreated exponentially growing cell culture) and negative control (1 mL of cells from the same culture heated to 80 °C for 5 min) were included with each run to gate the live and dead cell populations.

### Flow Cytometry

Cell abundances, viability (Live/Dead violet staining and chlorophyll fluorescence), and fluorescein-labelled oligonucleotide delivery were measured on a Guava easyCyte 12HT flow cytometer (EMD Millipore, Billerica, MA). Cells were excited with a blue 488 nm laser for measuring chlorophyll fluorescence (692/40 nm), size (forward scatter) and fluorescein fluorescence (525/30 nm); and with a violet 405 nm laser to measure the violet dead stain fluorescence (448/50 nm). All flow cytometry files were analyzed using FlowJo software version 7.6.5 (FlowJo, LLC, Ashland, OR, USA) and the following workflow: (1) *Prochlorococcus* cells were counted by a defined gate on the red fluorescence (chlorophyll) versus forward scatter (size) plot. (2) From these selected events, viable cells were counted by a defined threshold on the blue fluorescence (violet dead cell stain). The threshold was adjusted for each run using the positive (untreated cells) and negative (heat-shocked cells) control. (3) From the ‘viable cell’ subset of events, the cells into which DNA was delivered were counted by a defined threshold on the green fluorescence (fluorescein-labelled oligonucleotides). The threshold was adjusted so that <1% of events from the negative control (cells that did not enter into contact with the oligonucleotides) were included. The dead cell control comprised cells killed by heat treatment at 80 °C for 5 min. The flow cytometry plots were generated using FlowJo.

### Plasmids and mini-Tn5 transposon construction

We used plasmid mini-Tn5 pBAMD1-4 [60] to construct transposon tools for insertion into *Prochlorococcus* genomes. Genes encoding the GFPmutII and YFP reporters, shown to fluoresce efficiently in a wide range of cyanobacteria [56] were cloned by Gibson assembly behind the strong *Prochlorococcus* P *ccmk* promoter for Rubisco protein [95], well conserved in various *Prochlorococcus* and *Synechococcus* strains [86]. The pBAMD1-4 plasmid was linearized by digestion with HindIII-HF and EcoRI-HF (New England Biolabs # R3104S and # R3101S) at 37 °C for 2 h in Cutsmart buffer. The GFPmutII-encoding fragment was amplified from plasmid pCV0001 DNA using primers RL017 and RL018; the YFP-encoding fragment was amplified from plasmid pCV0021 using primers RL020 and RL021. The P ccmk promoter was amplified from *Prochlorococcus* MED4 genomic DNA using primers RL015 and RL016 for the P ccmk-gfpmutII construct, and RL015 and RL019 for the P ccmk-yfp construct. The 3-part assembly reaction (plasmid-promoter-insert) was performed using the Gibson assembly master mix 2x (New England Biolabs #E2611S), following the manufacturer’s instructions. The mixture was transformed in One Shot™ PIR1 Chemically Competent *E. coli* (Thermofisher #C101010). For the construction of pBAMD1-4-P ccmk (comprising the P ccmk promoter inserted within the transposon in front of the antibiotic resistance *aadA* gene, but no fluorescent marker gene) P ccmk was amplified from *Prochlorococcus* MED4 genomic DNA using primers RL121 and RL122, which contain EcoRI and SwaI restriction sites, respectively. Both pBAMD1-4 and the insert were digested with EcoRI-HF and SwaI-HF (New England Biolabs #R0604S and # R3101S), ligated, and transformed in PIR1 *E. coli* as described above.

The chloramphenicol resistance gene *cat* and the spectinomycin/streptomycin resistance gene *aadA* were codon-optimized for *Prochlorococcus* using the codon usage table from the Kazusa database, and the OPTIMIZER online tool [96]. The codon-optimized sequence, which is preceded by the P *ccmk* promoter and flanked by the Tn5 mosaic end, and were synthesized through the Genscript gene synthesis service (see Genbank sequence file as supplementary data).

### Transposome preparation

Transposome DNA containing the antibiotic resistance cassette and the mosaic ends were amplified using the template plasmids and primers (all 5’ phosphorylated) listed in table 2:

- Transposome T1 amplified using RL005 and RL006 primers from pCV0001 template DNA
- Transposome T2 amplified using RL007 and RL008 primers from pCV0021 template DNA
- Transposome T3 amplified using RL079 and RL080 primers from pBAMD1-4-GFP template DNA
- Transposome T4 amplified using RL079 and RL080 primers from pBAMD1-4-YFP template DNA
- Transposome T5 amplified using RL081 and RL080 primers from the synthesized codon-optimized chloramphenicol resistance cassette template DNA
- Transposome T7 amplified using RL079 and RL080 primers from the pBAMD1-4-P *ccmk* template DNA
- Transposome T5 amplified using RL140 and RL080 primers from the synthesized codon-optimized Spectinomycin/streptomycin resistance cassette template DNA.

PCR mixtures were digested for 1 h at 37 °C with DpnI (New England Biolabs) to remove the template plasmids. PCR products were then purified using the QIAquick PCR Purification Kit (Qiagen), eluted in TE buffer at pH 7.5, and DNA purity was assessed using a NanoDrop ND8000 spectrophotometer. Using the same kit, samples were often purified a second time to remove detectable impurities. Transposomes were formed using the EZ-Tn5™ Transposase (Lucigen) following the manufacturer’s instruction. 4 μL of EZ-Tn5 transposase preparation was mixed with 2 μL of transposon DNA at 100 μg μl^−1^ plus 2 μL of 100% glycerol and the mixture was incubated for 30 min at room temperature. The transposomes were either electroporated immediately or frozen at −20 °C for later use. The commercial transposome R6K*γ*ori/KAN-2 from Lucigen kit #TSM08KR was used according to the manufacturer’s instructions.

### Pour plating of axenic *Prochlorococcus* cultures

To obtain single colonies from an axenic culture, serial dilutions of an exponentially growing cyanobacterial culture grown in Pro99 medium (as described above) were pour-plated in the plating medium (Pro99 medium supplemented with 0.05% (wt/vol) Pyruvate and 3.75 mM TAPS (pH 8) and containing 0.3% low melting point (LMP) agarose. To prepare the plating medium, we first sterilized a 3% LMP agar solution in ultrapure water independently by three rounds of boiling in a microwave (~20 sec each, to avoid boiling-over and loss of volume). We then added the melted LMP agar 1:10 in the Pro99 medium base supplemented with pyruvate and TAPS. The mixture was maintained at ~28 °C in a water bath to cool before plating. Of importance, the LMP agarose was microwave sterilized and not autoclaved to avoid creating reactive oxygen species likely to inhibit *Prochlorococcus* growth [77]. Microwave-based boiling was sufficient to prevent the appearance of contaminants in the nutrient-poor Pro99 medium. For antibiotic selection after transposome delivery, 1 mL of ‘recovering’ cells (post electroporation, see above) were centrifuged and resuspended in 100 μL of fresh Pro99, and pour plated in 10 mL plating medium supplemented with the antibiotic. Kanamycin was used at 50 μg ml^−1^, and aminoglycoside resistance (encoded by *aadA*) selected with a combination of streptomycin and spectinomycin at 5 μg ml^−1^ total concentration (2.5 μg ml^−1^ each). Individual colonies typically became visible after 40-60 days.

They were picked using a sterile pipet tip and inoculated into 5 mL of fresh Pro99 medium. The efficiency of colony formation from a diluted culture is variable and generally low (~1%), an effect that seems alleviated by plating a higher number of cells, as an axenic lawn of cells would grow significantly faster, typically visible after 15-30 days.

## ACKNOWLEDGMENTS

We gratefully acknowledge Benjamin M. Woolston for sharing his method for DNA delivery screening in live cells, which was critical to the development of the procedure in *Prochlorococcus*.

This study was supported in part by the National Science Foundation (NSF-EDGE – 1645061 to SWC) and the Simons Foundation (Life Sciences Project Award IDs 337262 and 509034SCFY17 and SCOPE Award ID 329108, to SWC).

