## Supplementary Data for "Toward a genetic system in the marine cyanobacterium *Prochlorococcus*"

### SUPPLEMENTAL INFORMATION

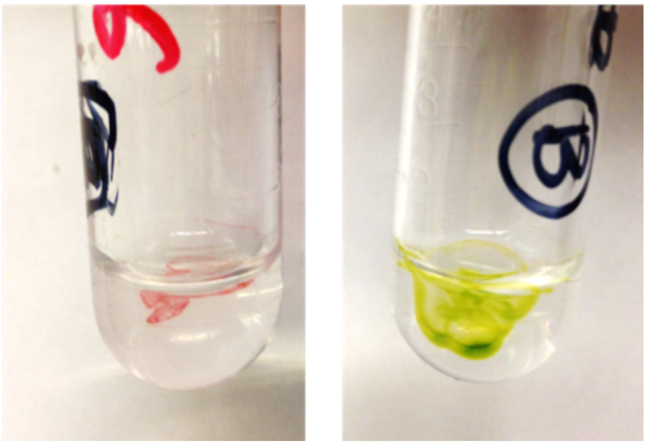

**Supplementary Fig. 1.** The 'agar stab' mating procedure. The picture shows the 1 mL agar stab following injection with 100 µL mixture of concentrated *E. coli* donor and receiver strain (*Synechococcus* strain WH7803 on the left, *Prochlorococcus* strain MIT9313 on the right). The tubes were placed in the constant light incubator for 24 h to allow mating (see methods for details).

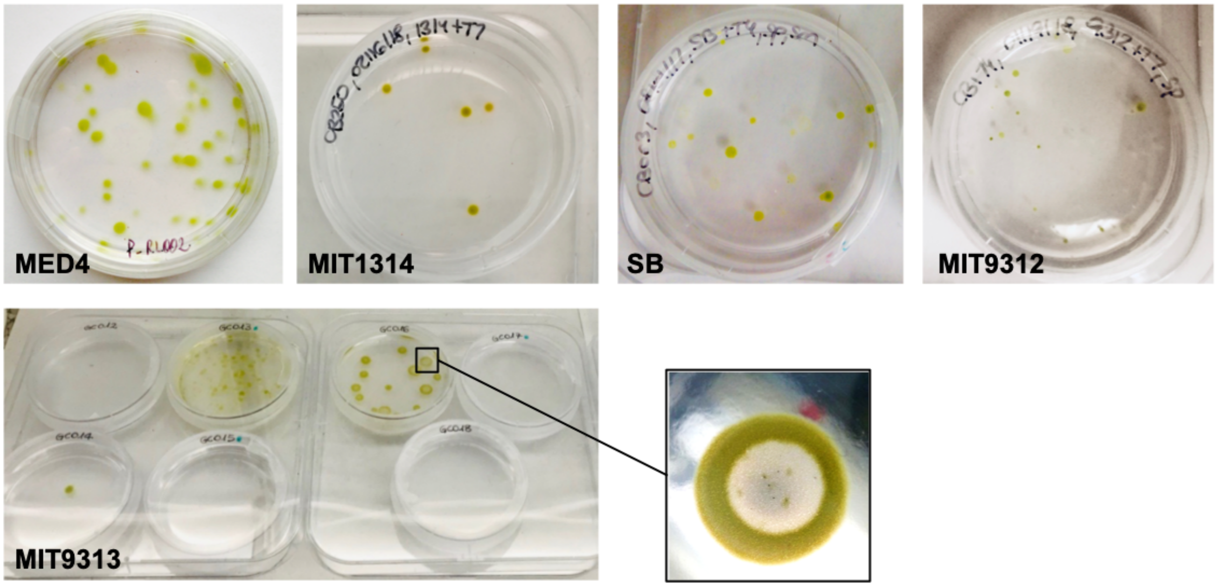


**Supplementary Fig. 2.** Example of axenic colonies obtained with various strains of *Prochlorococcus*. For all strains, colonies were incubated for at least six weeks. Colonies can appear more or less 'diffused' according to the strain and the incubation time. All colonies were growing within the agar and not at the surface. After eight weeks of growth, colonies sometimes adopt a ring shape, as cells in the center of the colony are dying (see close up picture of a ring-shaped colony).

**Supplementary File 1.** Transposome T5 sequence (codon-optimized chloramphenicol resistance).

**Supplementary File 2.** Transposome T7 sequence (codon-optimized streptomycin/spectinomycin resistance).
